# *Ex vivo* mycobacterial growth inhibition assay (MGIA) for tuberculosis vaccine testing - a protocol for mouse splenocytes

**DOI:** 10.1101/020560

**Authors:** Andrea Zelmer, Rachel Tanner, Elena Stylianou, Sheldon Morris, Angelo Izzo, Ann Williams, Sally Sharpe, Ilaria Pepponi, Barry Walker, David A. Hokey, Helen McShane, Michael Brennan, Helen Fletcher

## Abstract

The testing of vaccines for tuberculosis is costly and time-consuming, and dependent on preclinical animal challenge models and clinical trials. We have recently developed a mycobacterial growth inhibition assay (MGIA) to test vaccine efficacy *ex vivo*. This assay measures the summative effect of the host immune response and may serve as a novel tool to facilitate vaccine testing. It has generated much interest recently, and to facilitate technology transfer and reproducibility between laboratories, we here describe a detailed protocol for an *ex vivo* MGIA in mouse splenocytes.

## Introduction

Tuberculosis (TB) is a major global health problem, and new vaccines are urgently needed. With a lack of easily measurable correlates of protection, vaccine efficacy can currently only be measured in pre-clinical animal challenge models and clinical trials, both of which are time consuming and expensive.

We have recently developed an *ex vivo* mycobacterial growth inhibition assay (MGIA) to facilitate vaccine testing. This assay is a refinement of protocols first described by Marsay et al. and Kolibab et al.^1,2^ Our assay provides a direct read-out of the ability of a vaccine-induced immune response to control mycobacterial growth. Its *ex vivo* nature requires the use of animals; however, it has the potential to be a more rapid and cost-effective pre-clinical method than those currently used, and animal welfare is improved by cutting out the need for animal challenge experiments, which are time-consuming, expensive, and require specialist containment facilities. Furthermore, the assay could be used in future as an additional read-out in clinical trials.

To facilitate and encourage technology transfer between laboratories and vaccine developers in the TB vaccine field, we here describe a protocol for an *ex vivo* MGIA using mouse splenocytes in sufficient detail to enable researchers to establish this assay in their own laboratories. This protocol as presented here works reliably in our hands but has not been validated across laboratories or species. We invite other researchers who are working or would like to work with this assay to discuss the method and communicate their experience via bioRxiv comments in order to further optimise it and evaluate its reproducibility.

Please note, data discussed here are unpublished at the time of deposition of this article, and this article will be updated once published data are available. A summary of key results is given where relevant.

## ***Ex vivo* MGIA Methodology**

### Principle

The MGIA principle is based on the notion that host cells will inhibit mycobacterial growth in cell culture *ex vivo* if an effective, specific immune response is induced. Using total splenocytes from mice ensures that all necessary immune cells are present in near-physiological numbers and activation stages. Co-culture of mycobacteria with these cells *ex vivo* allows us to measure overall change in bacterial numbers, e.g. a reduction induced by vaccination. Bacterial numbers can be quantified using a BACTEC MGIT system, which assesses bacterial growth automatically every hour. This system is based on fluorescence quenching by oxygen - the silica gel at the bottom of the tube is non-fluorescent in the presence of the oxygen-enriched mycobacterial growth medium; however when bacteria grow and their metabolism reduces oxygen concentration, fluorescence becomes more intense. Once fluorescence reaches a certain threshold, the tube is registered as positive, and the time to detection (TTD) is given. TTD is dependent on the initial number of bacteria used to inoculate the MGIT tube, and the relationship between initial bacterial numbers and TTD can be determined by producing a standard curve as described below (Fig. 1). The standard curve can then be used to convert TTD to bacterial numbers (CFU; colony forming units).

**Fig. 1:**
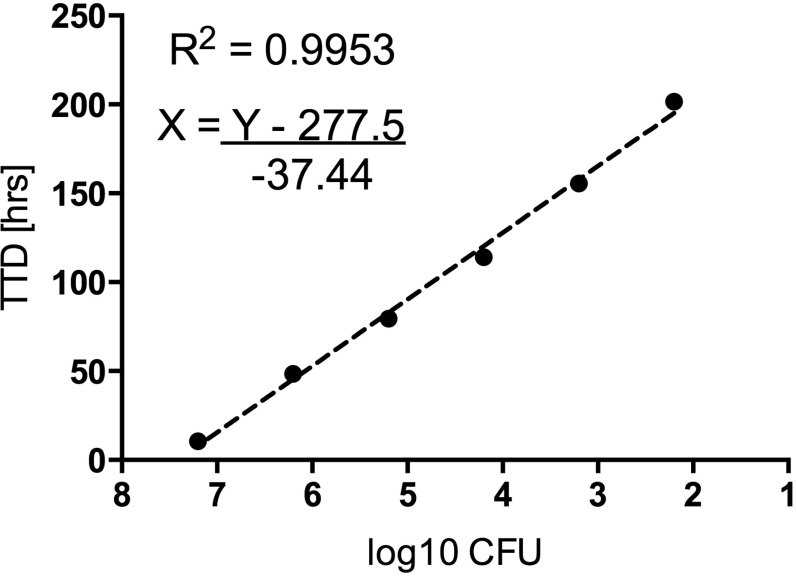
BACTEC MGIT standard curve. Total CFU were calculated from the number of colonies counted from plating the bacterial stock. CFU were converted to log10 CFU, and linear regression analysis carried out by fitting a semi-log log line within GraphPad Prism. The R^2^ value indicates the goodness of fit.

### Experimental design and expected outcomes

In our experience, total host cell numbers, as well as mycobacterial strain and inoculum dose, influence the difference found between vaccinated and naïve groups in this assay. We therefore recommend titrating both in a pilot experiment (Fig. 2).

In our hands and using the protocol described below, we found that using higher numbers of splenocytes (5×10^6^) produced greater differences between naive and BCG-vaccinated mice, and reduced variability within groups. An input dose of ∼50-100 CFU BCG Pasteur Aeras (which typically resulted in a TTD of approx. 8.5 days for the direct-to-MGIT controls) resulted in a reduction of 0.7-0.8 log10 CFU in splenocytes from vaccinated mice, a difference that was replicated in *Mycobacterium tuberculosis* challenge experiments. These results were obtained using C57Bl/6J mice that were immunised at the age of 6-8 weeks, and *M. bovis* BCG Pasteur, which was produced by the Aeras Laboratories (Rockville, MD) and is an early passage strain. We have found statistical power to be sufficient with group sizes of 6-8 animals.

★**NOTE**: The TTD for the input inoculum may vary depending on the mycobacterial strain used and a standard curve should be produced for each strain.

**Fig. 2:**
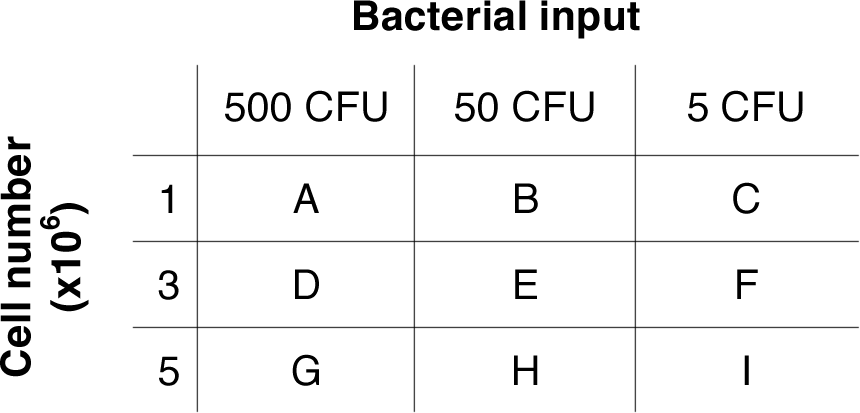
Outline of a potential titration experiment. Each cell number (1, 3, or 5 ×10^6^ splenocytes) is incubated with each bacterial input dose (500, 50, or 5 CFU). This results in 9 conditions and therefore 9 samples per animal (or 18 samples if prepared in duplicate).

### Procedures

The following sections give details on procedures as used in our laboratory. Standard materials (e.g. consumables such as tips or small reaction tubes) are used. Specific materials, which should not be substituted, are indicated in bold.

★**NOTE**: Each laboratory must ensure they are complying with local biological safety and ethics policies and procedures.

#### STANDARD CURVE

<u>Materials:</u>
- *M. bovis* BCG stock
- Sterile 1.5-2 ml microcentrifuge tubes
- Sterile filter tips 1000*μ*l, 200*μ*l and 20*μ*l
- **Barcoded BACTEC MGIT tubes (Cat. No. 245122, BD)**
- **PANTA enrichment (Cat. No. 245124, BD)**
- Sterile PBS + 0.05% Tween 80 (other diluent can be used as desired, such as water or growth medium from supplemented BACTEC MGIT tubes)
- 7H11 agar plates + 0.5% glycerol + 10% OADC
- sealable bags or parafilm

#### MGIT tube preparation - work under sterile conditions

1. Prepare MGIT PANTA enrichment: reconstitute one bottle of lyophilized MGIT PANTA by adding contents of one bottle MGIT growth supplement. Mix by turning over until completely dissolved.
2. Add 800 *μ*l of MGIT PANTA enrichment to each BACTEC MGIT tube. Prepare 2 tubes per dilution. Tightly recap the tube(s). Unused PANTA enrichment can be stored at +4°C for up to 1 month.

★**NOTE**: Keep tubes sealed whenever possible as they are oxygen-enriched.
★**NOTE**: Do not store MGIT tubes after the addition of PANTA enrichment. The tubes must be used the same day or discarded.

*Stock titration*

1. Thaw one vial of the mycobacteria to be tested at room temperature. **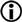OPTIONAL**: If the mycobacterial stock appears clumpy, it can be sonicated in a sonication water bath for 1 min. Place vial on ice for 1 min and repeat. ★**NOTE**: The homogeneity of the mycobacterial stock has an impact on the reproducibility of the results in this assay. It is important that the input dose is reproducible between experiments. Sonication might help to achieve this. If the stock preparation is homogenous and input doses are consistent without sonication, this step can be omitted.
2. Prepare 7 10-fold serial dilutions of the stock: add 1.08 ml PBS-Tween (or other diluent as desired) to each of 7 sterile microcentrifuge tubes.
3. Add 120 *μ*l undiluted stock to the first tube. Mix thoroughly by pipetting up and down.
4. From the first tube (1:10 dilution), remove 120 *μ*l and transfer it to the next tube, mix. Repeat until all 7 dilutions are prepared. ★**NOTE**: The number and range of dilutions can be adjusted according to the stock concentration, if known.
5. Inoculate 2 MGIT tubes for each dilution by adding 500 *μ*l of the appropriate dilution to each tube. Tightly seal tubes as soon as possible. Turn over to mix.
6. Place MGIT tubes into the BACTEC MGIT instrument. Note time to detection (TTD) as tubes come positive. ★**NOTE**: Record tube numbers (given on barcode) or position within the drawer for each sample – this will make it easier to identify samples from the MGIT report (Fig. 3).
7. Divide two 7H11 agar plates into 3-4 sectors each. Spot 3×20 *μ*l from each dilution onto a section. Leave plates to dry. Seal plates in sealable bags or with Parafilm to prevent drying out. Place in incubator at 37°C. Count colonies as soon as they are visible (after 3-4 weeks for most standard BCG strains; colonies of BCG Pasteur from Aeras are countable after approx. 10-12 days). Counts from spots should be approx. between 4 and 30 colonies.
8. Calculate standard curve by plotting TTD against input CFU as determined by plating or the equivalent volume (step 9) and use regression analysis to obtain the equation that can be used to convert any TTD to volume or CFU (Fig. 1; also see data analysis section). ★**NOTE**: TTD values for the standard curve should be between 1-20 days approx. ★**NOTE**: Software programmes such as Microsoft Excel or GraphPad Prism can be used for linear regression analysis.

#### IMMUNISATION

<u>Materials:</u>
- *M. bovis* BCG stock
- sterile disposable 1 ml syringe
- sterile 25G needle
- sterile saline
- sterile centrifuge tube (such as 50 ml Falcon tube)
- 7H11 agar plate

1. For immunisation with BCG, prepare inoculum by adding appropriate BCG stock volume to sterile saline to make a final concentration of 4×10^5^ CFU in 100 *μ*l saline per animal. Prepare/administer other vaccine candidates as appropriate.
2. Inject 100 *μ*l subcutaneously into each mouse (or other volume as per local guide lines).
3. Carry out the *ex vivo* MGIA as described below at the time point of the peak immune response after immunisation. ★**NOTE**: Our data were obtained at 6 weeks after the immunisation with BCG Pasteur Aeras in C57Bl/6J mice. However, the *ex vivo* MGIA should be carried out at the time point of the peak immune response, which is depending on vaccine candidate and mouse strain used.

#### *EX VIVO* MGIA

<u>Materials (all sterile except ice):</u>
- Ice
- 50ml falcon tubes
- Cell culture grade PBS
- Dissection kit
- Cell strainers 100 *μ*m
- Small petri dishes (approx. 50mm in diameter/10mm high)
- 5 ml disposable syringes
- Pastettes (disposable plastic Pasteur-type pipettes)
- RPMI-MGIT (w/o antibiotics): RPMI-1640 HEPES modification + 2 mM L-Glutamine + 10% FBS
- ACK buffer (NH_4_Cl (final concentration 150 mM), KHCO_3_ (final concentration 10 mM), Na_2_EDTA (final concentration 100 mM)), make up in milli-Q H_2_O, adjust pH to 7.2-7.4, autoclave); alternatively Sigma RBC lysis buffer or other lysis buffer as desired can be used.
- Cell culture grade water
- **MGIT tubes and PANTA enrichment, as above**
- **2ml screw cap tubes (Cat. no. 72.694.006, Sarstedt)**
- **Tube rotator (Cat. no. 444-0502, VWR UK)**
- 7H11 plates + 10% OADC + 0.5% Glycerol

**DAY 1** *Processing of spleens from mice*

★**NOTE:** Keep cells on ice whenever possible.
★**NOTE**: As cells will be co-cultured with bacteria, cells must be processed and cultured in antibiotic-free medium. Strict sterile working technique must be observed to avoid contamination.

1. Dissect spleens from mice aseptically, place in 5 ml sterile RPMI-MGIT medium in a 50 ml Falcon tube.
2. Pour spleen and 5 ml RPMI-MGIT into a cell strainer in a small petri dish.
3. Mash spleen through strainer by applying gentle pressure using the rubber end of a plunger from a 5 ml syringe.
4. Transfer cells back to 50 ml Falcon using a disposable pastette.
5. Centrifuge cells at approx. 450g for 5 min at 4°C.
6. Discard supernatant by decanting, loosen pellet by tapping the bottom of the tube, and add 5 ml Sigma RBC lysis or ACK buffer for approx. 2 minutes at room temperature.
7. Add 25 ml RPMI-MGIT (approx. 5x original volume) to dilute lysis buffer.
8. Centrifuge cells as above and discard supernatant.
9. Resuspend cells in 10 ml RPMI-MGIT and count viable cells using your standard method.
10. Adjust cell numbers to required concentration (e.g. 1, 3, or 5×10^6^ cells) per 300 *μ*l by adding RPMI-MGIT, prepare sufficient total volume for the number of samples needed (e.g. 9 × 300 *μ*l samples if using the above titration design).

*Preparation of mycobacterial mastermix*

1. Calculate total volume of bacterial suspension and concentration needed for all samples. Eg. if using the above titration design, per each animal prepare 3 × 300 *μ*l at 500 CFU/300 *μ*l, 3 × 300 *μ*l at 50 CFU/300 *μ*l, and 3 × 300 *μ*l at 5 CFU/300 *μ*l. For each bacterial concentration, include 2× 300*μ*l for direct-to-MGIT controls and excess for plating on 7H11 agar plates.
2. Thaw BCG stock at room temperature. **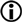OPTIONAL**: Sonicate stock vials using a sonicating water bath. This might not be necessary if stocks are homogeneous and no clumping is observed (also see note under step 3). Sonicate for 1 min, place on ice for 1 min. Repeat.
3. Add appropriate volume of stock to RPMI-MGIT to make up correct bacterial number per 300 *μ*l and sufficient total volume for all samples.
4. Vortex to mix.
5. Supplement 2 MGIT tubes for each BCG concentration with 800 *μ*l PANTA enrichment.
6. Add 300 *μ*l of each concentration of mycobacterial mastermix to each of 2 MGIT tubes. Add another 200 *μ*l of RPMI-MGIT to each MGIT tube to make the total volume added to the tube 500 *μ*l as per manufacturer’s recommendation. ★**NOTE**: These two duplicate tubes are the “direct-to-MGIT” positive controls. They serve as quality control to ensure reproducibility of input inocula, and can also be used to normalise data across experiments (see data analysis section). **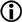OPTIONAL**: If using a live vaccine such as BCG, it may be desirable to set up control MGIT tubes to detect any residual BCG from the immunisation by adding 5 million cells in 300 *μ*l (from previously prepared cell suspension) from each mouse to a supplemented MGIT tube. Add 200 *μ*l RPMI-MGIT to make the total volume added to the tube 500 *μ*l.
7. Plate appropriate dilution of mycobacterial master mix on 7H11 plates. Spot 3x20 *μ*l drops for each dilution. Alternatively, if the concentration of the inoculum is very low, spread larger volumes (e.g. 50 *μ*l or 100 *μ*l) onto the plate.

*Cell/mycobacteria co-culture and assessment of growth inhibition*

1. Place 300 *μ*l of splenocytes at your desired concentration (e.g. 1x, 3x, or 5×10^6^ cells) into a labelled 2 ml screw cap tube.
2. Add 300 *μ*l mycobacterial mastermix of the appropriate concentration to each tube. **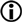OPTIONAL**: Samples can be set up in duplicate if desired; this is advisable at least for the first experiment to assess the quality of the technical replicates.
3. Place all 2 ml tubes on a 360° tube rotator inside a 37°C incubator for 96 hrs (4 days). Angle the paddle attachment at approx. 45°C to the middle axis.

**DAY 4** *Cell lysis and preparation of MGIT samples*

1. Supplement a MGIT tube for each sample with 800 *μ*l PANTA enrichment, and label. ★**NOTE**: Keep tubes sealed as much as possible as they are oxygen-enriched. ★**NOTE**: Do not store MGIT tubes after the addition of PANTA enrichment. The tubes must be used the same day or discarded.
2. Centrifuge the splenocyte/mycobacteria samples in a bench top microcentrifuge at 12,000 rpm for 10 min.
3. Remove 500 *μ*l of supernatant, ensuring the pellet remains intact. ★**NOTE**: Supernatants can be transferred to a new sterile 2 ml tube and stored at -20°C for further analysis if needed.
4. Add 400 *μ*l sterile tissue culture grade water to each tube.
5. Vortex each tube briefly.
6. Leave for 5 min at room temperature, vortex again.
7. Leave a further 5 min at room temperature, vortex again.
8. Add the total volume of each sample (now approx. 500 *μ*l) to the appropriate MGIT tube, invert to mix, and place in BACTEC MGIT machine.
9. Note TTD values for all samples and calculate CFU values based on the MGIT standard curve prepared previously (see data analysis section).

## DATA ANALYSIS

The previously prepared standard curve (Fig. 1) allows for conversion of the TTD given on the MGIT report (see Fig. 3 below for an example) to CFU. GraphPad Prism software works well for regression analysis and determining the equation that describes the relationship between the initial CFU input and TTD. Microsoft Excel provides a similar functionality, but we found Prism to provide more options.

★**NOTE**: Depending on whether CFU or log10 converted CFU values are used in the standard curve, the regression analysis is slightly different. Log10 CFU as shown in the example in Fig. 1 can be fitted with a linear regression, whilst using CFU requires the fitting of a semi-log line.

The regression analysis will provide you with a R^2^ value (a measure of how well the line fits the data), and the equation describing the line. Solve the equation for X: Y= A*X + B —> X = (Y-B)/A (where A = slope; B = intersect). By inserting the TTD (=Y), the number of CFU (or log10 CFU) initially added to the tube can now be calculated.

★**NOTE**: It is important to distinguish between total CFU and CFU/ml, as the TTD will indicate total numbers of bacteria per MGIT tube (i.e. in this assay, the number of bacteria in each MGIA sample (steps 30 & 31) of originally 600 *μ*l).

This data can now be plotted in a graph as is best suited for the experiment. If desirable, log10 CFU can be converted to CFU and vice versa. If more than one experiment is to be compared, it may be advisable to normalise data to the direct-to-MGIT controls to account for differences in input inocula. In this case, the number of total CFU per sample would be divided by the number of total CFU in the direct-to-MGIT control to give a read-out of relative growth, or fold-change in bacterial number.

**Fig. 3:**
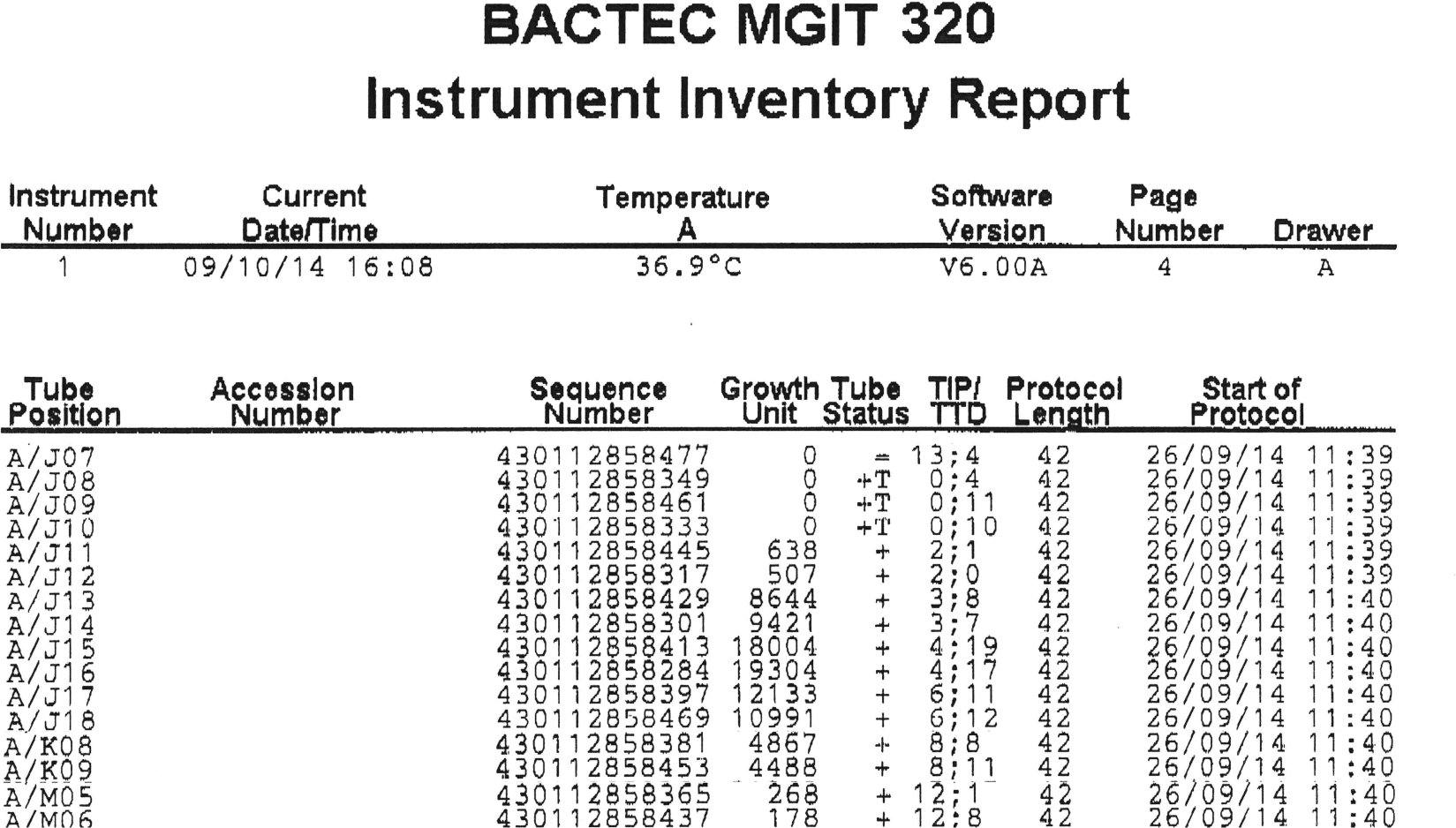
Example of a MGIT report (composit image). This figure shows the raw data for the standard curve displayed in Fig. 1. The report provides the following information:

Tube position: The position of the sample within the instrument, where the first letter indicates the drawer (if more than one), the second letter the row within the drawer, and the number the position within the row. These are clearly labeled within the instrument. Sequence number: The unique sequence number of the barcoded MGIT tube. Growth units: an expression of the measured fluorescence intensity. Tube status: + positive; - negative; = sample still in protocol. TIP/TTD: Time in protocol - the time the sample has been under investigation in the instrument; TTD - time to detection. TTD is given in the format “days;hours”. Protocol Length: this indicates for how long the sample will be monitored until declared negative (in days). This can be up to 56 days. Start of protocol: the date and time the sample is registered in the instrument; this field will be empty until the first reading is taken.

## Discussion

At a time when STOP TB and the WHO have called for an ambitious goal to end TB as a global epidemic by 2035 (an average less than 10 tuberculosis cases per 100,000 population; cut the number of deaths from TB by 95%), funding to achieve this goal grows more scarce and it is now more important than ever for scientists within the TB community (and other fields) to join forces, use the available budgets strategically, and to seek new avenues for vaccine testing and development. This is reflected by the encouragement of funding bodies to form large research consortia such as TBVAC2020 and EMI-TB. In an effort to facilitate technology transfer and help standardise vaccine testing, between and within these groups, we describe here a detailed protocol for an *ex vivo* Mycobacterial Growth Inhibition Assay (MGIA) which offers an alternative approach to currently used cytokine-based assays for the assessment of new TB vaccine candidates. ^3,4^

These assays measure directly the summative ability of host cells to control mycobacterial growth, a more holistic assessment criterion for novel vaccine candidates than the *in vitro* assays currently used. These assays also provide a unique opportunity to manipulate the host response in a very defined manner and measure the effect on the ability to control bacterial growth, e.g. by neutralisation of cytokines, depletion of specific host cell populations, or use of genetically modified strains of both animals or mycobacteria. Although similar assays have been described^1,2,5,6^, they have not been widely adopted by TB vaccine developers as they have been perceived as difficult to establish and lacking in reproducibility. This manuscript provides detailed descriptions of both laboratory methods and data analysis for a pre-clinical *ex vivo* MGIA method first described by Marsay et al. We aim to facilitate the uptake of this assay by TB vaccine developers. We detail the critical considerations for establishment of this assay and provide an assay optimisation strategy for transfer of the assay into a new laboratory and assessment of a new mycobacterial infection stock.

For this assay to be a useful tool for vaccine assessment in humans, blood is the only feasible sample to use, and a PBMC-based, as well as a whole blood assay are currently under development.^7,8^

It may be possible to determine bacterial numbers at the end of the 4 day incubation period by conventional plating of the lysed cells on 7H11 agar plates; however, we have not yet tested this approach. It would make these experiments more cost effective, and independent of the availability of a BACTEC MGIT system. On the other hand, the BACTEC MGIT system provides a wide dynamic range and sensitivity in the determination of the bacterial load that is not matched by culture on solid media, where several dilutions of one sample may have to be plated.

All in all we believe that this assay will be of use for many groups within the TB research community, and will contribute to the development of an effective vaccine against tuberculosis.

## References

1. Marsay, L. et al. Mycobacterial growth inhibition in murine splenocytes as a surrogate for protection against Mycobacterium tuberculosis (M. tb). Tuberculosis (Edinb). 93, 551–7 (2013).

2. Kolibab, K., Yang, a., Parra, M., Derrick, S. C. & Morris, S. L. Time to detection of *Mycobacterium tuberculosis* using the MGIT 320 system correlates with colony counting in preclinical testing of new vaccines. Clin. Vaccine Immunol. 21, 453–455 (2014).

3. Soares, A. P. et al. Bacillus Calmette-Guérin vaccination of human newborns induces T cells with complex cytokine and phenotypic profiles. J. Immunol. 180, 3569–77 (2008).

4. Beveridge, N. E. R. et al. A comparison of IFNgamma detection methods used in tuberculosis vaccine trials. Tuberculosis (Edinb). 88, 631–40 (2008).

5. Kolibab, K. et al. A practical *in vitro* growth inhibition assay for the evaluation of TB vaccines. Vaccine 28, 317–22 (2009).

6. Parra, M. et al. Development of a murine mycobacterial growth inhibition assay for evaluating vaccines against *Mycobacterium tuberculosis*. Clin. Vaccine Immunol. 16, 1025–32 (2009).

7. Fletcher, H. A. et al. Inhibition of mycobacterial growth in vitro following primary but not secondary vaccination with *Mycobacterium bovis* BCG. Clin. Vaccine Immunol. 20, 1683–9 (2013).

8. Burl, S., Holder, B. S., Lo, B. K. M. & Kampmann, B. Optimisation of a functional mycobacterial growth-inhibition assay to improve its suitability for infant TB vaccine studies. J. Immunol. Methods 394, 121–4 (2013).

